# A new colorectal cancer risk prediction model incorporating family history, personal and environmental factors

**DOI:** 10.1101/662106

**Authors:** Yingye Zheng, Xinwei Hua, Aung K. Win, Robert J. MacInnis, Steven Gallinger, Loic Le Marchand, Noralane M. Lindor, John A. Baron, John L. Hopper, James G. Dowty, Antonis C. Antoniou, Jiayin Zheng, Mark A. Jenkins, Polly A. Newcomb

## Abstract

**Purpose:** Reducing colorectal cancer (CRC) incidence and mortality through early detection would improve efficacy if targeted. A CRC risk-prediction model incorporating personal, family, genetic and environmental risk factors could enhance prediction.

**Methods:** We developed risk-prediction models using population-based CRC cases (N=4,445) and controls (N=3,967) recruited by the Colon Cancer Family Registry Cohort (CCFRC). A familial risk profile (FRP) was calculated to summarize individuals’ risk based on their CRC family history, family structure, germline mutation probability in major susceptibility genes, and a polygenic component. Using logistic regression, we developed risk models including individuals’ FRP or a binary CRC family-history (FH), and risk factors collected at recruitment. Model validation used follow-up data for population-(N=12,052) and clinic-based (N=5,584) relatives with no cancer history at recruitment, assessing calibration (E/O) and discrimination (AUC).

**Results:** The E/O (95% confidence interval [CI]) for FRP models for population-based relatives were 1.04 (0.74-1.45) and 0.86 (0.64-1.20) for men and women, and for clinic-based relatives 1.15 (0.87-1.58) and 1.04 (0.76-1.45). The age-adjusted AUC (95% CI) for FRP models in population-based relatives were 0.69 (0.60-0.78) and 0.70 (0.62-0.77), and for clinic-based relatives 0.77 (0.69-0.84) and 0.68 (0.60-0.76). The incremental values of AUC (95% CI) for FRP over FH models for population-based relatives were 0.08 (0.01-0.15) and 0.10 (0.04-0.16), and for clinic-based relatives 0.11 (0.05-0.17) and 0.11 (0.06-0.17).

**Conclusion:** The FRP-based model and FH-based model calibrate well in both settings. The FRP-based model provided better risk-prediction and discrimination than the FH-based model. A detailed family history may be useful for targeted risk-based screening and clinical management.

## INTRODUCTION

Screening for colorectal cancer (CRC) is efficient and cost effective.^1^ The evidence is compelling, even when applied irrespective of personal characteristics except age.^2^ However, beyond age, individuals’ CRC risk factors could inform the use and frequency of specific screening regimens.^3,4^ A detailed risk prediction model would permit targeted screening at an appropriate level based on individuals’ risk of CRC.^5-8^

Family history of CRC is an important risk factor for this disease, as it is a proxy for genetic and environmental factors shared by relatives.^9,10^ A comprehensive risk prediction model would incorporate detailed family history of cancer and available information on known genetic, and epidemiologic characteristics. To date, existing CRC or colorectal adenoma risk prediction models are limited,^7^ including simple measures of CRC family history and limited risk factor data,^11-16^ or considering only a small group of known low-penetrance SNPs that explain little familial aggregation of CRC.^17^ Moreover, of the seven models that have been validated using external samples, they were found to have only reasonable discrimination, suggesting limited usefulness for risk-based screening.^18^

The majority of these risk models defined family history as a binary variable, typically as “at least one first-degree relative with CRC”; a few models considered family history as the number of first- or second-degree relatives with CRC. A small number of models (e.g., MMRpro)^19^ used a more complex definition based on the number of affected relatives, their ages of CRC diagnosis and the degree of relatedness. In theory, the more detailed and accurate the family history, the better the risk prediction. However, in a typical primary care setting with limited time and incomplete patient reports, only the presence or absence of a CRC family history is generally recorded. It is unclear whether such information is sufficient to predict CRC risk accurately. Using the large well-characterized population-based data from the Colon Cancer Family Registry Cohort (CCFRC), ^20,21^ we describe the development and validation of a new risk prediction model that incorporates a novel measure of family history in addition to personal and environmental risk factors.

## METHODS

### Study sample

The CCFRC is an NCI-funded international consortium of six CRC registries from the USA, Canada, and Australia/New Zealand, using standard protocols for data collection, molecular characterization, and follow-up at each site (http://coloncfr.org/). Recently diagnosed CRC cases from population-based cancer registries, controls from population-based sources (including drivers’ license, voting records, health beneficiary rosters, and electoral rolls), and cases from family cancer clinics with a strong family history of CRC or early-onset disease were recruited as “probands” between 1998-2012.^20^ Relatives of population- and clinic-based cases were also invited to participate. Informed consent was obtained from participants in all study sites. Local institutional research ethics review boards approved the study protocols.

### Data collection and testing

At baseline, participants were asked to: (i) complete an epidemiological risk factor questionnaire on medical history, demographic characteristics, reproductive history, physical activity, medication, postmenopausal hormone use, alcohol and tobacco use, and diet about one year before diagnosis or a comparable period in controls; (ii) describe detailed CRC family history information, at least for their first degree relatives, including relationship to the participant, age, sex, and type and ages of cancer diagnosis; (iii) provide written consent for the research team to access tumor tissues and corresponding pathological reports; and (iv) collect a blood or buccal sample. Reported cancers and ages at diagnosis were confirmed, where possible, using pathology reports, medical records, cancer registry reports and/or death certificates. Genetic mutation screening and testing for mismatch repair (MMR) and *MUTYH* mutations was completed for probands and relatives, as previously described in detail.^22^ During follow-up approximately every 5 years, participants from all case families were contacted for updates on incident polyps, cancer diagnoses at any site, CRC screening and surgery, as well as among their relatives cancer diagnoses and deaths. Population-based controls were not followed up.

### Family history measures

Family history of CRC was defined in two ways: 1) as a binary indicator (yes/no) of having at least one first-degree relative with CRC (hereafter called, FH); and 2) as a continuous familial risk profile (FRP) based on detailed cancer family history, considering age at disease diagnosis as well as the number of and relationship to each relative, their ages, and their probabilities of carrying CRC predisposing mutations in the DNA MMR genes (*MLH1, MSH2, MSH6, PMS2*) and *MUTYH* genes.^22,23^ The FRP can be considered as a probability index, indicating absolute risk of CRC from birth to age 80 years. The FRP was calculated based on modified segregation analysis using: (i) the age- and sex-specific incidence of CRC from national cancer statistics;^24^ (ii) the familial relative risk based on previous segregation analysis of CRC data from the CCFR;^22^ and (iii) the age-specific incidence of CRC based on mutation status, for which we used the penetrance reported from analysis of the CCFRC (**Supplementary Methods**).^22,25-29^ We included established pre-diagnosis risk factors in model development, including screening (as described in **Tables 1-2** and **Supplementary Methods**). A list of candidate variables and their parameterization used for model selection are described in **Supplementary Table 1**.

**Table 1.** Associations between risk factor variables and coloretal cancer from the risk model with familial risk profile (FRP model) and from the risk model with a binary family history (FH model), for men only

| Variables | Cases (N= 2312)* | Controls (N= 1916)* | FRP Model† | FH Model† |
| --- | --- | --- | --- | --- |
|  | No. (%) | No. (%) | OR (95% CI) | OR (95% CI) |
| Family History |  |  |  |  |
| FRP, Mean (SD) | 0.09 (0.097) | 0.07 (0.027) | 1.16 (1.11, 1.20)<sup>¶</sup> | -- |
| Binary FH, No. (%) |  |  |  |  |
| No | 1859 (80.4) | 1731 (90.3) | -- | 1.00 (Ref) |
| Yes | 453 (19.6) | 185 (9.7) |  | 2.34 (1.90, 2.88) |
| Recent BMI<sup>§</sup>, kg/m<sup>2</sup> |  |  |  |  |
| <25 | 571 (24.6) | 600 (31.3) | 1.00 (Ref) | 1.00 (Ref) |
| 25-30 | 1113 (48.0) | 940 (49.1) | 1.38 (1.18, 1.62) | 1.35 (1.15, 1.58) |
| >30 | 611 (26.3) | 372 (19.4) | 1.59 (1.31, 1.93) | 1.61 (1.32, 1.95) |
| Red meat consumption, servings/day |  |  |  |  |
| <1 | 1681 (72.4) | 1484 (77.5) | 1.00 (Ref) | 1.00 (Ref) |
| 1+ | 564 (24.3) | 364 (19.0) | 1.27 (1.08, 1.51) | 1.25 (1.06, 1.47) |
| Regular NSAID use duration<sup>II</sup>,<br>years |  |  |  |  |
| Non-user | 1364 (58.8) | 939 (49.0) | 1.00 (Ref) | 1.00 (Ref) |
| ≤ 2 | 475 (20.5) | 388 (20.3) | 0.91 (0.76, 1.09) | 0.93 (0.78, 1.10) |
| > 2 | 406 (17.5) | 509 (26.6) | 0.73 (0.61, 0.87) | 0.72 (0.61, 0.86) |
| Calcium supplement use duration,<br>years |  |  |  |  |
| Non-user | 2060 (88.8) | 1610 (84.0) | 1.00 (Ref) | 1.00 (Ref) |
| ≤ 2.5 | 115 (5.0) | 97 (5.1) | 1.29 (0.95, 1.75) | 1.32 (0.97, 1.79) |
| > 2.5 | 84 (3.6) | 137 (7.2) | 0.62 (0.46, 0.83) | 0.62 (0.46, 0.83) |
| Cigarette Smoking, pack-years |  |  |  |  |
| Never | 810 (34.9) | 689 (36.0) | 1.00 (Ref) | 1.00 (Ref) |
| <10 | 332 (14.3) | 281 (14.7) | 1.00 (0.81, 1.23) | 1.03 (0.84, 1.27) |
| 10 to 19 | 289 (12.5) | 229 (12.0) | 1.18 (0.95, 1.47) | 1.16 (0.93, 1.45) |
| 20+ | 807 (34.8) | 632 (33.0) | 1.31 (1.11, 1.55) | 1.34 (1.14, 1.58) |
| History of polyp‡ |  |  |  |  |
| No | 2079 (89.6) | 1691 (88.3) | 1.00 (Ref) | 1.00 (Ref) |
| Yes | 159 (6.9) | 197 (10.3) | 1.46 (1.10, 1.95) | 1.46 (1.10, 1.94) |
| History of FOBT‡ |  |  |  |  |
| No | 1569 (67.6) | 1175 (61.3) | 1.00 (Ref) | 1.00 (Ref) |
| Yes | 646 (27.8) | 667 (34.8) | 0.79 (0.67, 0.93) | 0.80 (0.68, 0.95) |
| History of sigmoidoscopy‡ |  |  |  |  |
| No | 1893 (81.6) | 1397 (72.9) | 1.00 (Ref) | 1.00 (Ref) |
| Yes | 330 (14.2) | 434 (22.7) | 0.86 (0.71, 1.05) | 0.85 (0.70, 1.03) |
| History of colonoscopy‡ |  |  |  |  |
| No | 2035 (87.7) | 1536 (80.2) | 1.00 (Ref) | 1.00 (Ref) |
| Yes | 217 (9.3) | 332 (17.3) | 0.47 (0.37, 0.60) | 0.48 (0.38, 0.61) |

BMI: Body mass index; FRP: Familial Risk Profile; FH: binary family history; FOBT: fecal occult blood test; NSAID:
Nonsteroidal anti-inflammatory drugs; OR= odds ratio; CI = confidence interval.
\* Do not sum to total due to missing values; N for each category of variables unless otherwise specified.
† Both models adjusted for study site and age at reference (age at diagnosis for cases and age at baseline interview for controls) in addition to the variables presented in this table.
‡ History of polyp, FOBT, sigmoidoscopy and colonoscopy were defined as history of having any of these conditions/test two years prior to enrollment.
§ As of two years before enrollment.
|| Regular NSAID use was defined as use of aspirin and/or ibuprofen at least twice a week for more than a month.
¶ OR for FRP: per 10% increase.

**Table 2.**
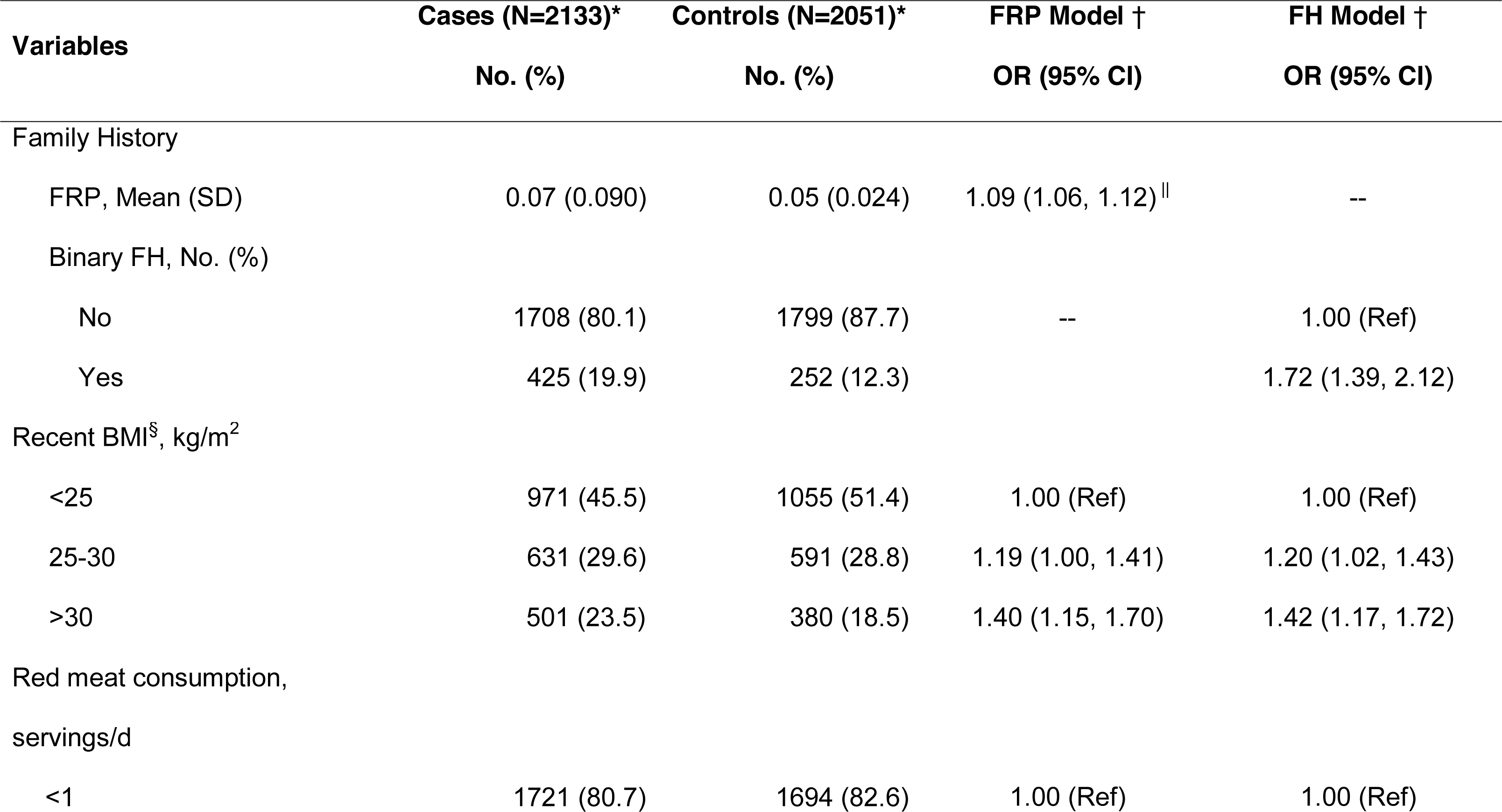
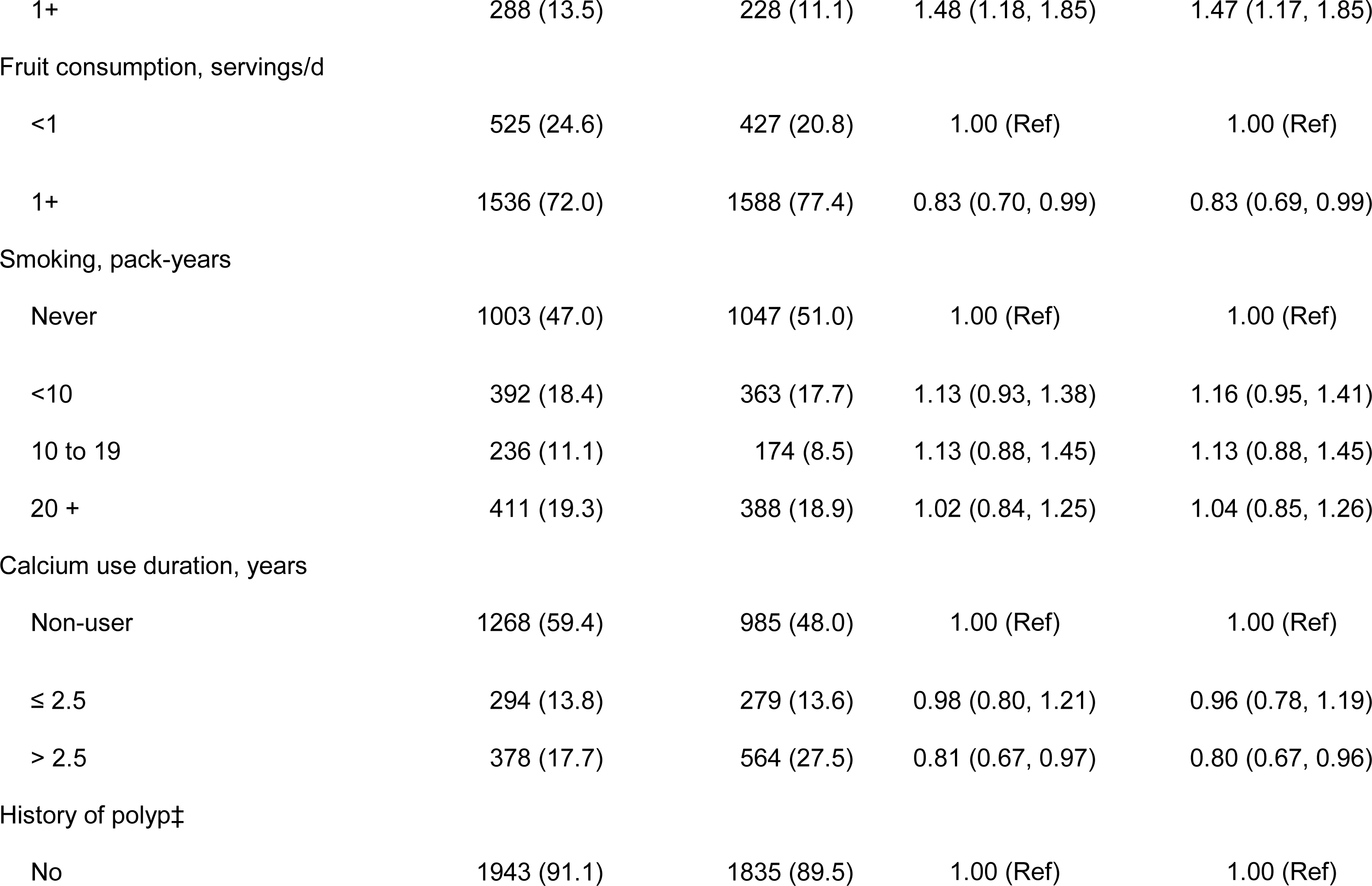
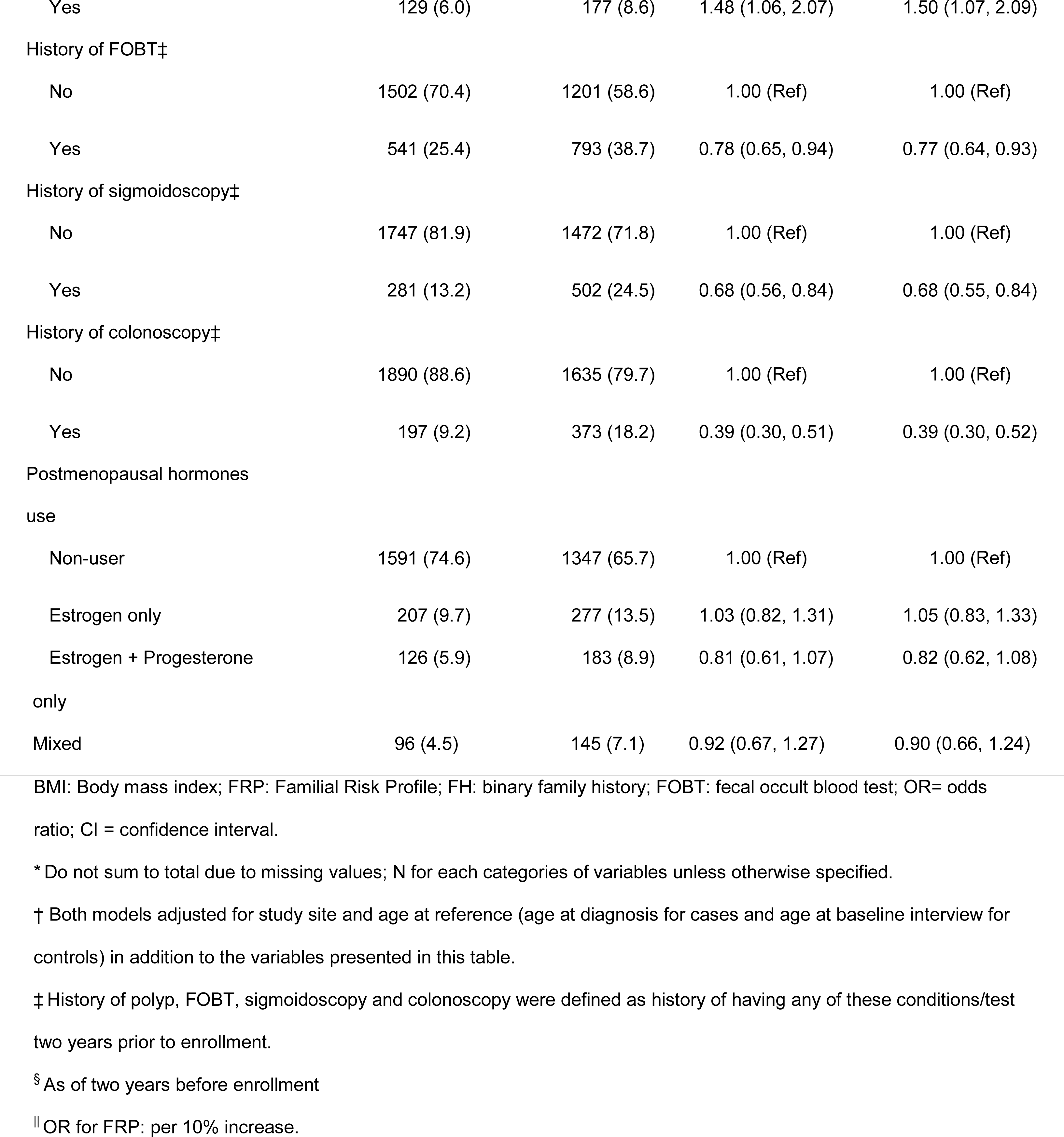
Associations between risk factor variables and colorectal cancer from the risk model with familial risk profile (FRP model) and from the risk model with a binary family history (FH model), for women only

## Statistical analysis

### Model development

We studied 4,445 CRC cases and 3,967 controls recruited from three population-based sites of the CCFRC (Seattle, WA, USA; Ontario, Canada; and Victoria, Australia). We only included cases who were diagnosed with CRC less than two years before completing the baseline questionnaire to ensure all risk factor pertained to the pre-diagnostic period. Controls were frequency-matched to the cases on age. We restricted our analysis to non-Hispanic whites. Participants with missing values on any of the baseline data variables were excluded from model development.

The CRC risk prediction model was developed using unconditional logistic regression with case/control status as the outcome. The distributions of FRP by sex and case-control status were examined using histograms and compared using Wilcoxon non-parametric tests. Models using either log-transformed FRP or FH were stratified by sex to permit sex-specific associations with risk factors. We adjusted for study site and reference age (age at diagnosis of CRC for cases and age at interview for controls) in all models. To account for site-specific sampling of CCFRC, we applied probability weights based on the (inverse) sampling probability of each individual’s selection into the study.^20^

Three forward-stepwise-selection procedures were implemented. Each used a different selection criteria: 1) *P* value < 0.15; 2) incremental value in Area Under the Receiver Operating Characteristic (ROC) Curve (incAUC) ≥ 0.01; or 3) a smaller Akaike information criterion (AIC).^30,31^ Final models from these different selection criteria were compared to identify variables that were robust to selection procedures. For example, if having a colorectal polyp was included in the variable set in a model based on incAUC, and as well as that from a AIC-based selection procedure, then it would then be included as a predictor in the final model. Two definitions of family history (FH and log-transformed FRP) were included in our final models in addition to the final list of environmental factors.

### Relative and absolute risk calculation

Odds ratios (ORs) and corresponding 95% confidence intervals (CI) were generated from the final models. To calculate absolute risk, we obtained sex- and age-specific CRC incidences for the USA (SEER-9 Registries, whites), Australia (Victoria) and Canada (Ontario) populations from the <u>Cancer Incidence in Five Continents</u> (CI5), for 1998-2002,^29^ corresponding approximately to the time period during which cases were diagnosed. Deaths from non-CRC causes were considered as competing risks. Age- and sex-specific mortality from causes other than CRC were obtained from all-cause mortality^32-34^ and CRC-specific mortality^35^ for the USA, Australia and Canada respectively during the same time period. Five-year absolute risks were calculated as described in Freedman et al.^14^ (**Supplementary Methods**).

### Model Validation

For model validation, we studied 17,636 unaffected relatives of case probands who: (1) were recruited from population registries (N=12,052) and genetic clinics (N=5,584); (2) were non-Hispanic whites; (3) had no personal history of any cancer at the time of recruitment; (4) completed a baseline questionnaire; and (5) were recruited from five study sites of the CCFRC (Australia/New Zealand, Mayo Clinic, Ontario, Cedars-Sinai and Seattle);^20^ and (6) were prospectively followed up after the baseline recruitment. The flowchart of the study design and model steps is included as **Supplementary Figure 1**.

We calculated absolute risk for each individual in the validation set based on our final model, the derived baseline risk functions, and data from the baseline questionnaire. In total, 12.6% of men and 17.0% of women had at least one covariate with missing data (see **Supplementary Table 2**). We conducted imputation for each risk factor variable in the final model with the most frequent (modal) category for men and women separately.^36^ Model performance was compared before and after exclusion of relatives with missing covariates (**Supplementary Table 3**). Using the follow-up data of the CCFRC, we identified incident CRC diagnoses. Time to event was defined as years from the date of baseline interview completion to the date of diagnosis of incident CRC. Individuals with a diagnosis of other types of cancer were censored at the date of diagnosis. Deaths due to causes other than CRC were considered as competing risk events. Participants who were alive without any cancer diagnosis were censored at the date of last contact.

Given different incidence rates for disease in the general population and the genetic clinic pool, we assessed model performance separately for population- and clinic-based relatives. For calibration performance, we categorized participants into quintiles of predicted absolute risk based on the developed model and plotted the average observed absolute risk within each quintile against the predicted risks in that quintile, adding 95% CIs of the observed risks. The observed marginal risks were calculated as the cumulative incidences of CRC accounting for censoring and competing risks.^37^ In addition, we calculated a summary measure of calibration for the FRP and FH models as the ratio of the averaged predicted 5-year absolute risk to the observed cumulative incidence rate (E/O), separately for men and women; 95% CIs were calculated using a bootstrap approach. ^38^ To assess the performance of the model for risk stratification, we defined four risk-groups based on the 30^th^, 60^th^ and 90^th^ percentiles of the predicted 5-year absolute risks of CRC. For each model, we plotted cumulative incidence functions of CRC diagnosis by risk-groups, as above, and tested differences among risk functions across groups using K-sample test.^39^ Since clinic-based population have higher CRC incidence rate than the rates from population-based registries, for clinic-based relatives, we calculated their baseline risk using the clinic-based set from our validation population.

For discrimination performance, we conducted Receiver Operating Characteristic (ROC) curve analyses and calculated AUC (**Supplementary Methods**) to assess the model’s ability to separate individuals with and without a CRC diagnosis within 5 years after baseline. Since the outcome of individuals who were censored within 5 years after baseline was uncertain, we excluded these individuals in this calculation and used inverse censoring probability weights to account for the missing information. Two age groups were defined as ≤50 and >50 years old at baseline separately for both men and women. Age-adjusted ROC curves and AUCs were calculated as the weighted average of age-specific AUC, with weights as the proportion of CRC diagnosis in each age group. We used a bootstrap approach to calculate the empirical 95% CIs for age-adjusted AUC based on 2.5 and 97.5 percentiles. Analyses were conducted using R version 3.1.1 (http://www.r-project.org/). All statistical tests were two-sided and *P* values of less than 0.05 were considered statistically significant.

## Results

### Model development

Using population-based cases and controls, the variables entered into the final models of FRP and FH for men and women are shown in **Tables 1 and 2**, respectively. The distribution of FRP (range: 0.037 to 0.993) by sex and case-control status is summarized in **Table 3** and **Supplementary Figure 2**. For both men and women, cases had higher FRP than controls (all *P* < 0.001).

**Table 3.** Distribution of Familial Risk Profile (FRP) by sex and case-control status.

| Sex | N | Mean (SD) | Minimum | Median | Q1-Q3 | Maximum | P value* |
| --- | --- | --- | --- | --- | --- | --- | --- |
| Men |  |  |  |  |  |  |  |
| Cases | 2312 | 0.091<br>(0.098) | 0.053 | 0.067 | 0.060-0.077 | 0.993 | <0.001 |
| Controls | 1916 | 0.068<br>(0.015) | 0.055 | 0.067 | 0.061-0.069 | 0.362 |  |
| Women |  |  |  |  |  |  |  |
| Cases | 2133 | 0.066<br>(0.091) | 0.037 | 0.045 | 0.043-0.052 | 0.979 | <0.001 |
| Controls | 2051 | 0.048<br>(0.023) | 0.038 | 0.045 | 0.043-0.047 | 0.917 |  |

\* P value for comparison of distributions between cases and controls was calculated using two-sided Wilcoxon non-parametric test; Q1: the first quartile; Q3: the third quartile

For men, every 10% relative increase in FRP (e.g. 0.33 vs 0.30) was associated with 16% higher risk of developing CRC (95% CI: 11%-20%). From the FH model, the OR for family history was 2.34 (95% CI: 1.90, 2.88). The strengths of association with the other variables were similar for FRP and FH models (**Table 1**). For women, a 10% relative increase in FRP was associated with 9% higher risk of CRC (95% CI: 6%-12%). From the FH model, the OR for family history was 1.72 (95% CI: 1.39-2.12). The strengths of associations with other variables were essentially no different than those from the FRP model (**Table 2**).

### Model validation

The median follow-up time was 8.6 years; 317 relatives were diagnosed with incident CRC during this period. Calibration for population- and clinic-based relatives across a wide range of risk groups is presented in **Supplementary Figure 3**. The overall E/O estimates (95% CI) for different models are summarized in **Table 4**. For population-based relatives, FRP and FH models calibrated well, with E/O estimates (95% CI) of 1.0 (0.7-1.4) and 0.9 (0.6-1.2) for men and women from FRP models, and (0.6-1.2) and 0.8 (0.6-1.2) from FH models. For clinic-based relatives, FRP and FH models calibrated well with E/O ranging from 1.0 to 1.2.

**Table 4.**
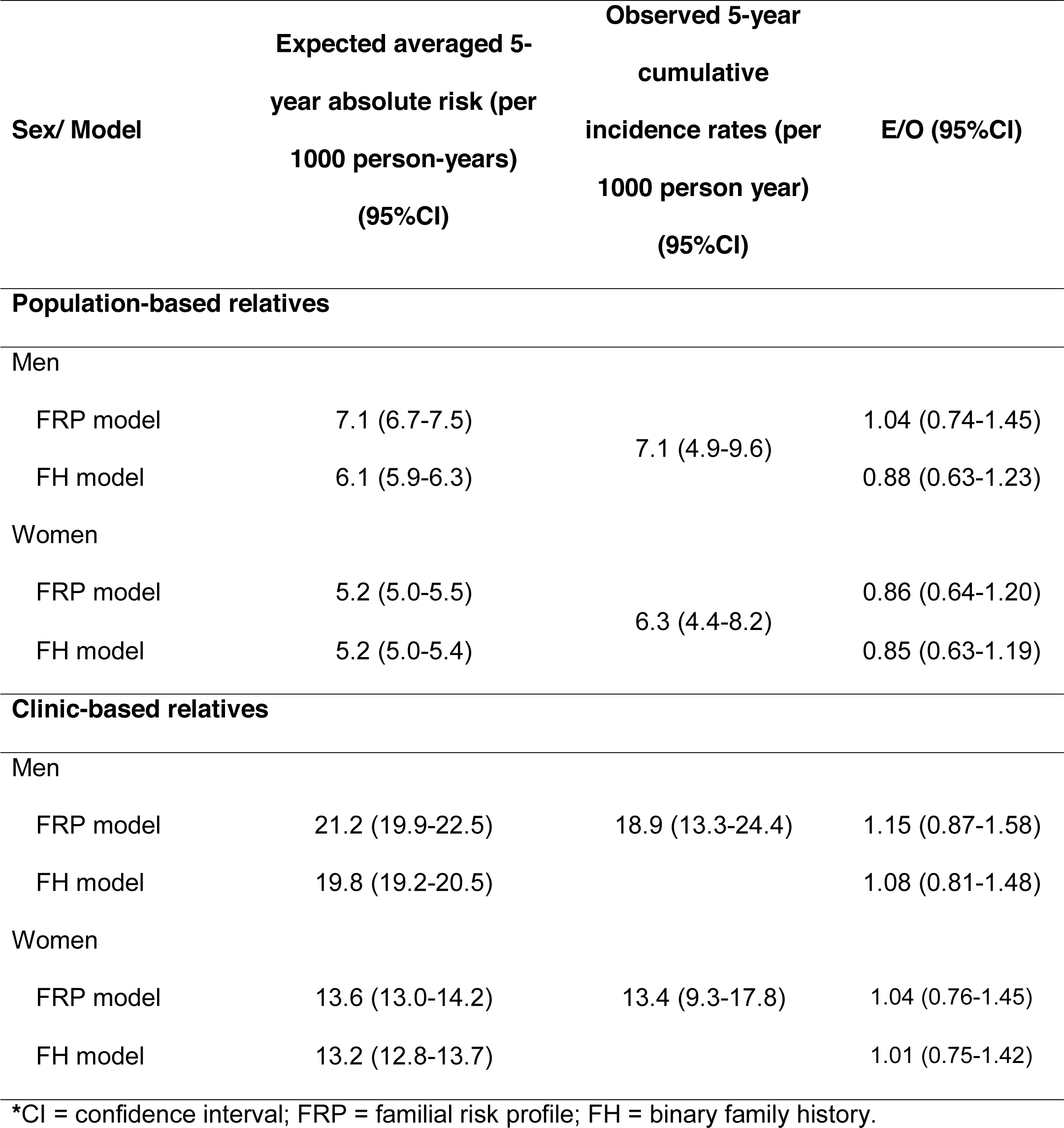
Observed 5-year cumulative incidence rates (O) versus Averaged 5-year absolute risk (E) based on risk models with familial risk profile or with a binary family history, and separately for men and women*

In addition, we defined four groups at different levels of predicted risks (using 30^th^, 60^th^, 90^th^ percentiles as cutoffs). The cumulative incidence curves are presented for population- and clinic-based relatives separately (**Figure 1a and 1b).** The wider separation of the FRP models suggests they performed better than the FH models in stratifying individuals into distinctive risk groups.

**Figure 1:**
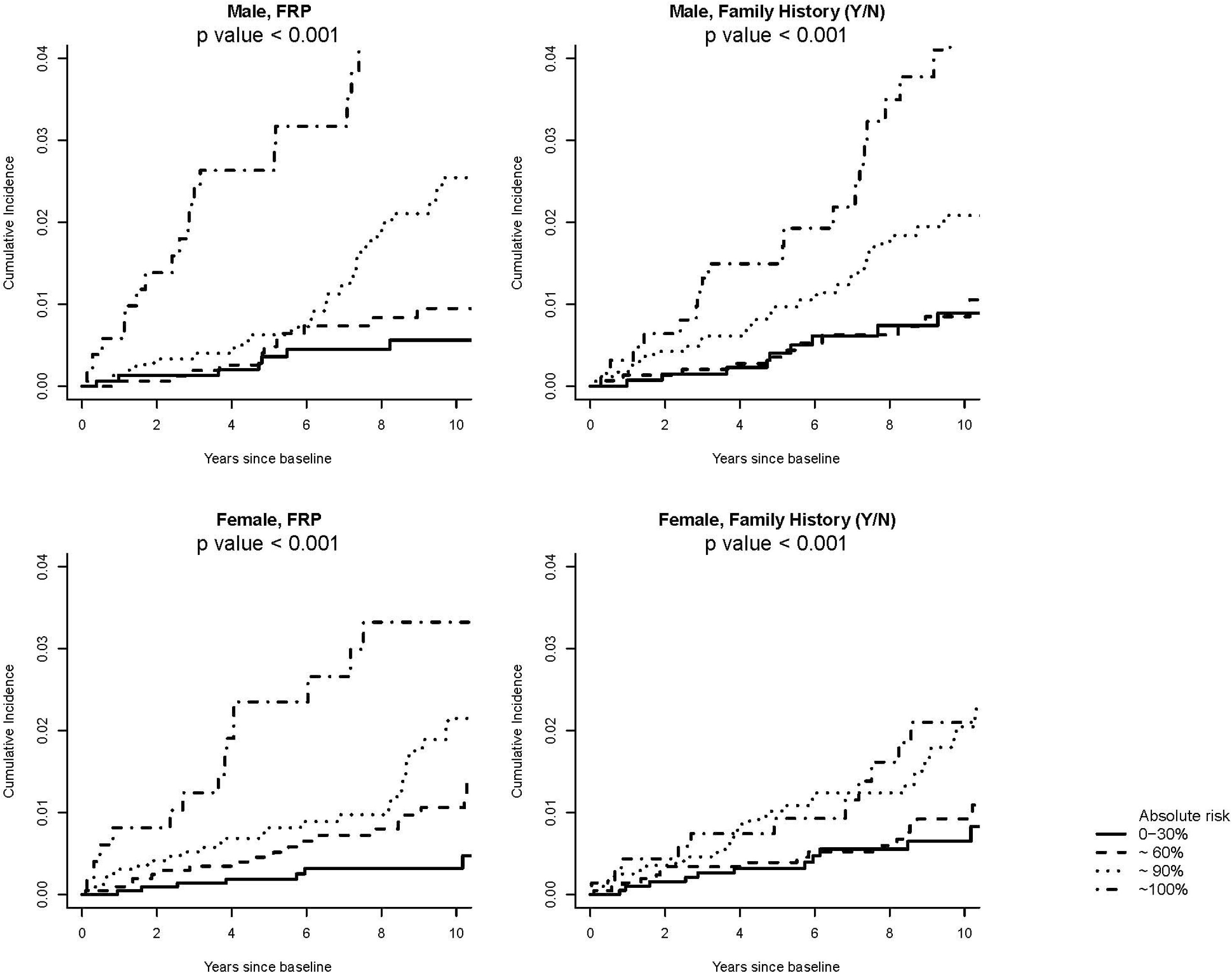
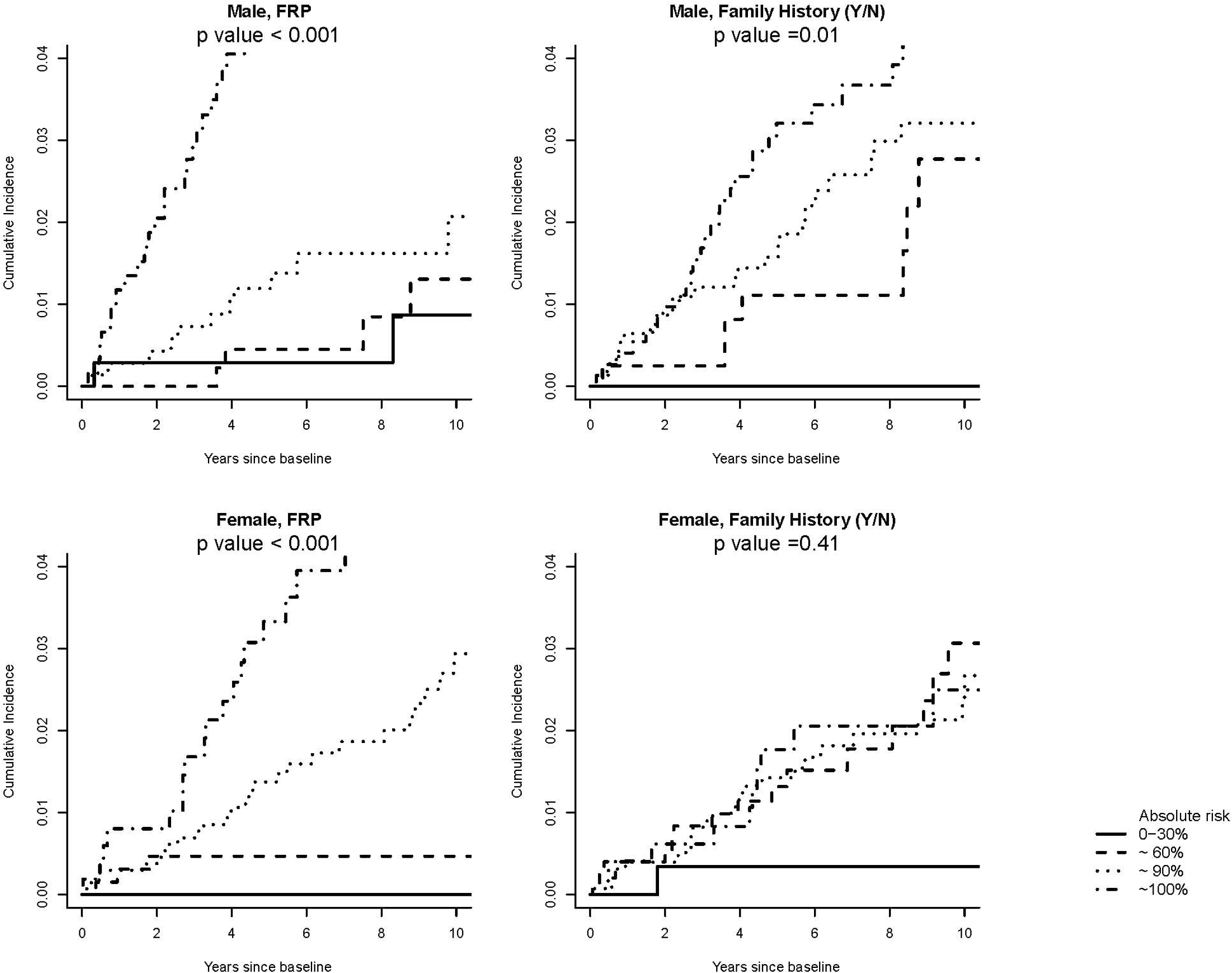
Cumulative incidence of colorectal cancer (CRC) according to estimated 5-year absolute risk among **a) population-based relatives, b) clinic-based relatives**. Four groups were defined based on cut-points of 30^th^, 60^th^ and 90^th^ percentiles of estimated 5-year absolute risk. The K-sample test was used to compare the cumulative incidence across groups and to calculate two-sided *P* values.^39^ FRP = familial risk profile

The FRP model also provided improved discriminatory capacity over the FH model (**Figure 2**). For population-based relatives, the age-adjusted AUCs for the FRP model was 0.73 (95%CI=0.67-0.79) for men and 0.70 (95%CI=0.62-0.77) for women. The increments in age-adjusted AUC (incAUC) for FRP over FH models were 0.08 (95%CI=0.01-0.15) for men, and 0.10 (95%CI=0.04-0.16) for women (both excluding the null with *P* < 0.001). For clinic-based relatives, the age-adjusted AUCs (95%CI) for FRP models were 0.77 (0.69-0.84) and 0.68 (0.60-0.76) for men and women, respectively. The incAUC (95% CI) for FRP over FH models was 0.11 (0.05-0.17) for men and 0.11 (0.06-0.17) for women.

**Figure 2.**
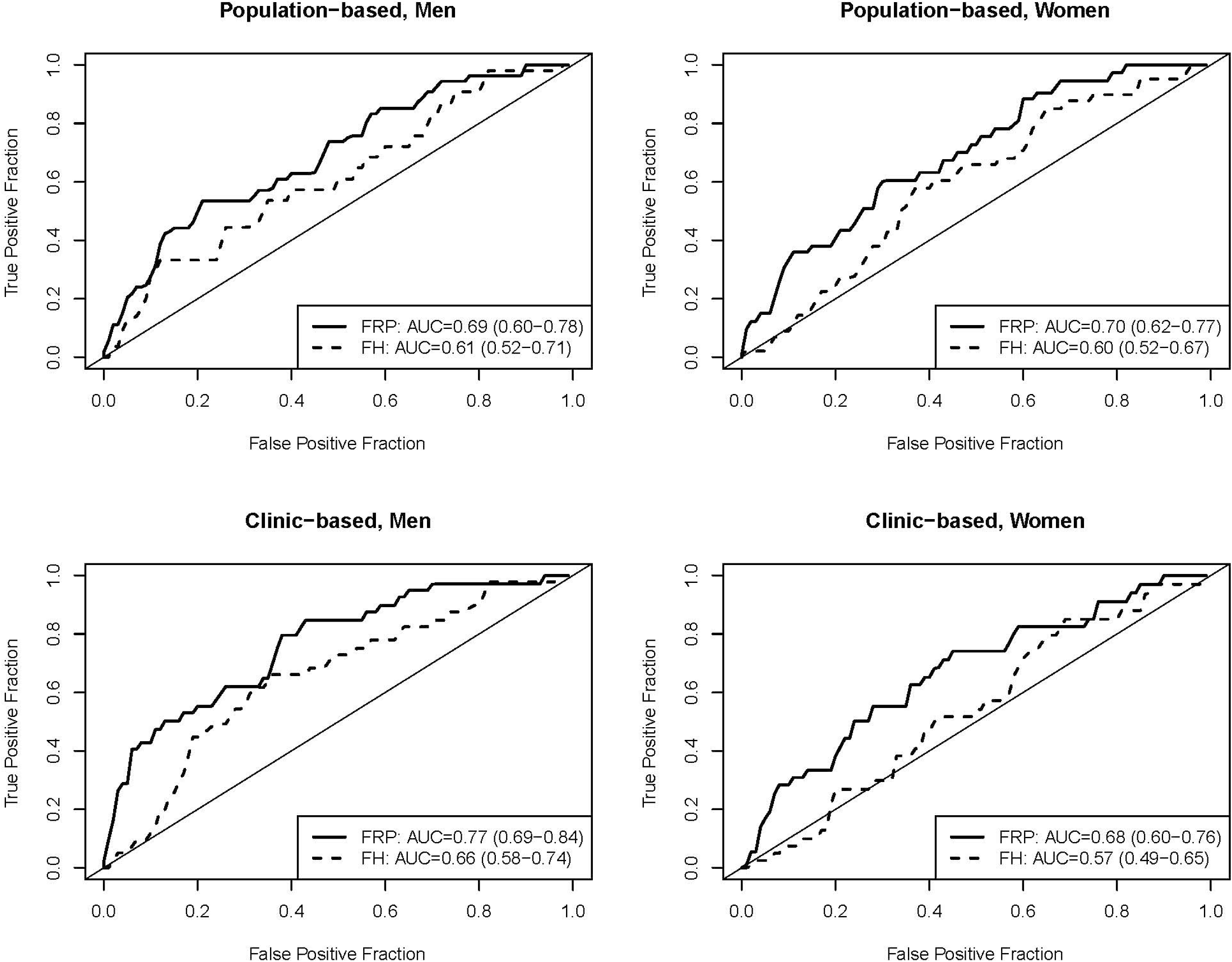
Age-adjusted Receiver Operating Characteristic (ROC) curves for men and women. ROC curves and Age-adjusted area under the curve (AUC) were calculated as the weighted average of age-specific estimates, with weights as the proportion of CRC diagnosis in each age group (<50 and >=50 at baseline). We calculated 95% confidence intervals (in parentheses) using bootstrap approach. FRP = familial risk profile; FH = binary family history.

## Discussion

We developed and validated a new risk prediction model which incorporated detailed family history information captured by the FRP, as well as personal and environmental risk factors. Generally, both FRP and FH models provided good calibration, however, our results also suggested that the FRP-based model gave better discrimination than a model using a simple binary summary of family history.^1^

One clinical utility of CRC risk models is to provide information for screening regimens tailored to an individual’s risk, and to inform intensity of screening, decisions regarding chemoprevention, and utilization of gene panel testing. Current CRC screening recommendations are based solely on age and simple measures of family history.^40^ Our study suggests that consideration of multiple risk factors, including a detailed family history of CRC, can lead to the identification of individuals across the spectrum of CRC risk, from those at very low risk with delayed and/or non-invasive screening recommendations, to those at high risk for whom earlier screening and more frequent/invasive monitoring is recommended. We have shown that family history of CRC is an important factor for CRC risk prediction, either defined as a binary (yes/no) measure or based on FRP calculated from the family structure, cancer histories, and MMR/MUTYH mutation status. Our research supports two approaches to risk prediction for CRC. In settings where family history information is limited, the risk model could include only the simple present/absent question. In settings such as genetic clinics where family history information is likely to be more complete, the risk model could make use of the FRP to derive more precise risk discrimination.

Numerous risk models have been developed to predict CRC and colorectal adenomas based on CRC family history, genetic mutation screening, personal characteristics, and known risk factors – singly or in combination.^7,18^ However, our FRP-based risk model is unique in its incorporation of all these CRC risk factors and in its use of our novel familial risk measure based on detailed family history information. Risk models that use family history as a binary indicator do not account for variability in family size, age, or structure, age of CRC diagnosis, or the relationship of affected relatives to the proband, which are integral to characterizing familial risk.^9,23,41^ Both models evaluated in our study included environmental factors, as have most prediction models, to take advantage of the substantial contribution of these exposures on CRC risk.^42^

Our study has many strengths, including its population-based design for the model development, and cases a broad spectrum of familial risk. All risk factors were collected by the CCFRC sites using the same instrument. In particular, the assessment of family structure and cancer history was extensive.^23,41^ Finally, the CCFRC’s use of a prospective follow-up design provided a validation data set with the same well-annotated information and from the same cohort upon which the model was developed.

Our study had some limitations. Although the validation of the model was prospective with epidemiologic factors assessed at baseline interview, the development of the model was based on retrospective reports of lifestyle and environmental exposures prior to recruitment. Additionally, since our cohort started over a decade ago, information on CRC screening might not reflect the most current screening practices, with newer screening tests and intervals now recommended.^40^ Further, since the absolute risks were derived based on the age-, sex-, and country-specific incidence from the general population, our model is well calibrated to population-based samples. We recalibrated the baseline risks for the clinical-specific model, using the clinic-based subset from our validation dataset. Future studies are needed to independently evaluate the calibration of this model especially for high-risk families. In addition, susceptibility SNPs identified by recent GWAS should be included to enhance the FRP-based model.^43,44^

In conclusion, we developed and validated a new CRC risk prediction model that incorporates a novel measure of family history, the FRP, in addition to personal characteristics and other non-family history-based risk factors. The new FRP-based model provided better risk discrimination than the FH-based model, suggesting that more detailed family history has the potential to be more informative for risk-based clinical decision-making.

## Acknowledgements

The authors thank all study participants of the Colon Cancer Family Registry and staff for their many contributions to this project.

## Footnotes

1 This model is available online at http://crisptool.org/crisp-int

## Notes

**Funding:** This work was supported by (R01 CA170122) from the National Cancer Institute, National Institutes of Health (NIH) and through cooperative agreements with the following Colon Cancer Family Registry (CCFR) centers: Australasian Colorectal Cancer Family Registry (U01/U24 CA097735), Mayo Clinic Cooperative Family Registry for Colon Cancer Studies (U01/U24 CA074800), Ontario Familial Colorectal Cancer Registry (U01/U24 CA074783), Seattle Colorectal Cancer Family Registry (U01/U24 CA074794), and USC Consortium Colorectal Cancer Family Registry (U01/U24 CA074799). Seattle CCFR research was also supported by the Cancer Surveillance System of the Fred Hutchinson Cancer Research Center, which was funded by Control Nos. N01-CN-67009 and N01-PC-35142 and Contract No. HHSN2612013000121 from the Surveillance, Epidemiology and End Results (SEER) Program of the National Cancer Institute. Additional support included grants from the National Institutes of Health UM1/U01 CA167551, K05 CA152715 (to PAN) and R01 GM085047 (to YZ), and through the Centre for Research Excellence grant APP1042021 and Program Grant APP1074383 from the National Health and Medical Research Council (NHMRC), Australia. MAJ is a NHMRC Senior Research Fellow. AKW is a NHMRC Early Career Fellow. JLH is a NHMRC Senior Principal Research Fellow.

The authors have no conflicts of interest to disclose.

