## Supplementary Materials for "A new colorectal cancer risk prediction model incorporating family history, personal and environmental factors"

**Supplementary Methods**

**A. Calculation of FRP**

The calculation involves two steps.

Step 1: Segregation Analysis

We used modified segregation analysis to fit a genetic model to the observed colorectal cancer family histories (age of onset) for the probands and *every* first- and second-degree relative. Individuals were assumed to be at risk of colorectal cancer from birth until the earliest of the following: diagnosis of colorectal cancer or any other cancer, first polypectomy, death, last known age at baseline interview or age 80 years. To model any residual familial aggregation of colorectal cancer risk, a mixed model that incorporated an unmeasured polygene in addition to the major gene was fit. Under this model, the colorectal cancer incidence for an individual *i* was assumed to follow the following parametric survival model

$$\lambda_{i}\left( t \right)=\lambda_{0}\left( t \right)\exp\left[ G_{i}+P_{i}\left( t \right) \right] \left( 1 \right),$$

where $\lambda_{0}\left( t \right)$ is the baseline incidence at age *t*. $G_{i}$ is the natural logarithm of the relative risk associated with the major genotype and was defined by six components representing each of the genes *MLH1*, *MSH2*, *MSH6*, *PMS2*, *MUTYH* and one representing the hypothetical unidentified major genes. The polygenic component for age *t*, $P_{i}\left( t \right)$, was assumed to be normally distributed with zero mean and variance $\sigma_{p}^{2}$. The hypergeometric polygenic model [1-3] was used to approximate *P*. Under Model (1), the colorectal cancer risk of individual *i* by age *t*, is

$P(T_{i} \leq t|$ $g_{i},p_{i}\left( u \right))=1-exp\left\{ -\int_{0}^{t} \lambda_{0}\left( u \right)\exp\left[ p_{i}\left( u \right) \right]du\exp\left( g_{i} \right) \right\} (2).$

Models were fit by maximum likelihood with the statistical package MENDEL version 3.2.6 [4]. Estimates were appropriately adjusted for the population-based ascertainment of families using a combination of retrospective likelihood and ascertainment corrected joint likelihood [1, 2] in which each pedigree’s data were conditioned on the proband’s genotype, cancer status and age of onset. Details of the models were reported in [5] and estimates of the model were reported in Table 3 of [5]. The FRP was calculated using the mixed dominant model in the table.

Step 2: Calculation of FRP

The Familial Risk Profile (FRP) is an estimate of the individual’s cumulative risk of colorectal cancer from birth to age 80 given their family history (FH), which includes their relative’s history of colorectal cancer (including ages at diagnoses) ***t***_R_ their personal mutation status of genes associated with colorectal cancer genetic syndromes (namely *MUTYH* and the DNA mismatch repair genes *MLH1, MSH2, MSH6,* and *PMS2*), *g*, and their relative’s mutation status of these genes, *g*_R_. Specifically, the FRP for individual *i* by age *80* is the probability of CRC diagnosis by age *80* given $g_{i},p_{i}\left( u \right),$ and age of onset of relatives $\boldsymbol{t}_{\boldsymbol{R}}$, which can be written as

$$P\left( T_{i}\leq80 \right| t_{R}, g_{i},p_{i}\left( u \right),g_{R})=\frac{P[T_{i}\leq80, t_{R} |g_{i},p_{i}\left( u \right), g_{R}]}{P[t_{R}|g_{R},p_{i}\left( u \right)]}$$

Hence, the risks can be computed as the ratio of two likelihoods. Specifically, the parameters needed to calculate this FRP are: (i) $\lambda_{0}\left( u \right):$the age- and sex-specific incidence of colorectal cancer in the relevant populations for the relevant years of the lives of the individuals, which we obtained from the relevant national cancer statistics sources for USA, Canada and Australia [6]; (ii) the familial relative risk based on their family history, which we obtained from a previous segregation analysis of colorectal cancer data from the Colon Cancer Family Registry [5]; and (iii) the age-specific incidence of colorectal cancer based on their mutation status, for which we used the penetrance reported from analysis of the Colon Cancer Family Registry and as specified in Equation (2) [5, 7-10]). Each of the likelihoods can be estimated using the formula provided in the supplement of [10]. The estimates of FRP for all individuals used for validation were obtained by the MENDEL package.

For individuals with unknown mutation status, their probability of being a mutation carrier was based on their family history of colorectal cancer using a previous segregation analysis [5] and, if available, their family history of a known mutation. This approach is the same used to estimate breast cancer risk based on family history and mutation status in *BRCA1* or *BRCA2* by the BOADICEA program [2].

**B. Calculation of Absolute Risk for Developing CRC**

The absolute risk for developing CRC in the age interval ($a_{0}$, $a_{0}$+$a$),$R\left( a | a_{0};Z \right)$, is the probability of developing CRC in a future time *a*, given that one is disease free at age $a_{0}$ and having a risk profile vector $Z$ at $a_{0}$. Let $A$ denote the age of onset of CRC, the absolute risk is defined mathematically by

$$R\left( a | a_{0};Z \right)=\Pr\left( a_{0}\leq A\leq\left( a_{0}+a \right) | T\geq a_{0},Z \right)=\int_{a_{0}}^{a_{0}+a} \lambda_{c}\left( u | Z \right)exp\left[ -\int_{a0}^{u} \left( \lambda_{c}\left( s | Z \right)+\lambda_{m}\left( s | Z \right) \right)ds \right]du,$$

where $\lambda_{c}\left( s | Z \right)$ is the hazard rate for CRC incidence, and $\lambda_{m}\left( s | Z \right)$ for competing causes of death other than CRC (see Supplemental Note of [11]). In addition, we assume $\lambda_{c}\left( s | Z \right)= \lambda_{c0}(s)\{rr\left( Z \right)\}$, where $rr\left( Z \right)=\exp\left( \beta Z \right),$ with $\exp\left( \beta\right)$ denoting the relative risk of $Z$ based on the logistic regression models. The age specific baseline hazard $\lambda_{c0}\left( s \right)$is calculated by $\lambda_{c0}\left( s \right)=\lambda_{c}^{*}\left( s \right)\times\left( 1-AR \right)$, where $\lambda_{c}^{*}\left( s \right)$ is the composite hazard rate based on the summation of proximal, distal, and rectal cancer incidence rates obtained from USA (SEER-9 Registries, whites), Australia (Victoria) and Canada (Ontario) populations from the Cancer Incidence in Five Continents (CI5), International Agency for Research on Cancer (IARC) from 1998 to 2002 [6]. In addition, AR denotes the attributable risk, and is estimated by

$$AR=1- \sum_{i=1}^{n_{D}} \frac{w_{i}Y_{i}}{rr\left( Z_{i} \right)},$$

with $w_{i}$denoting the weight accounting for sampling scheme used by the CCFR for the *i*th case ($Y_{i}=1)$, and summation is over all $n_{D}$ cases. In this manuscript, we also assume that $\lambda_{m}\left( s | Z \right)= \lambda_{m}\left( s \right),$and $\lambda_{m}\left( s \right)$can be similarly calculated using all-cause mortality and CRC-specific mortality for the USA, Australia and Canada respectively during the same time period.

All estimates were fit separately by sex.

**C. Calculation of ROC Curves under Censoring**

A validation cohort, consisting of healthy relatives of probands at baseline followed for CRC outcomes, is used for the ROC analysis. Denote T as the time from baseline to CRC diagnosis, and C as the censoring time defined as the time from baseline to time of last follow up or termination of the study. We define T* = min(T,C), and δ =1 if T* = T. To calculate the ROC curve for discriminating individuals who will develop CRC in the next 5-years (cases: T < 5 years) versus these who will remain disease free (controls: T>5 years), individuals who were censored before 5 years were dropped out from the analysis. To account for such incomplete data, we considered an inverse probability of censoring weighed (IPCW) approach [12, 13], in which individuals contributing to the estimation were inversely weighted by the probability of being included in the analysis. Specifically, at a risk threshold *p*, the time-dependent true and false positive fraction (TPF and FPF) can be estimated as

$${TPF}_{5 year}(p)= \frac{\sum_{i} w_{i}^{c}\delta_{i}I(T^{*}\leq5)I\left[ R\left( a_{i} | 5;Z_{i} \right)\geq p \right]}{\sum_{i} w_{i}^{c}\delta_{i}I(T^{*}\leq5)}$$

$${FPF}_{5 year}\left( p \right)= \frac{\sum_{i} w_{i}^{c}I\left( T^{*}>5 \right)I\left[ R\left( a_{i} | 5;Z_{i} \right)\geq p \right]}{\sum_{i} w_{i}^{c}I\left( T^{*}>5 \right)} ,$$

where the contribution of an observation needs to be weighted by

$$w_{i}^{c}\left( t=5 years \right)=\delta_{i}I\left( T^{*}\leq5 \right)\frac{1}{G\left( T_{i}^{*} \right)}+I\left( T^{*}>5 \right)\frac{1}{G(5)},$$

with G(s) = P(C > s) is the censoring probability and can be estimated using the Kaplan-Meier estimator. The 5-year ROC curve can be obtained by plotting ${TPF}_{5 year}(p)$ versus ${FPF}_{5 year}(p)$ across all *p*, and the AUC is

$${AUC}_{5 year}=\int{TPF}_{5 year}\left( p \right)d{FPF}_{5 year}\left( p \right).$$

**Supplementary Tables**

**Supplementary Table 1. List of candidate variables for model selection**

| **Candidate variables** | | **Total**  **(N=8412)** | **Male (N=4228)** | | **Female (N=4184)** | |
| --- | --- | --- | --- | --- | --- | --- |
|  |  |  | **No. of Cases (%)** | **No. of Controls (%)** | **No. of Cases (%)** | **No. of Controls (%)** |
| CRC Familial Risk Profile Score, Median (IQR) | | 0.059 (0.0230) | 0.0668 (0.0167) | 0.0667 (0.0082) | 0.0452 (0.0091) | 0.0445 (0.0040) |
| Family History of CRC in 1st degree relatives | |  |  |  |  |  |
|  | 0 | 7097 | 1859 (80.4) | 1731 (90.3) | 1708 (80.1) | 1799 (87.7) |
|  | ≥1 | 1315 | 453 (19.6) | 185 (9.7) | 425 (19.9) | 252 (12.3) |
| Age, years | |  |  |  |  |  |
|  | <50 | 2871 | 942 (40.7) | 451 (23.5) | 945 (44.3) | 533 (26) |
|  | ≥ 50 | 5487 | 1346 (58.2) | 1465 (76.5) | 1158 (54.3) | 1518 (74) |
|  | Unknown | 54 | 24 (1.0) | 0 | 30 (1.4) | 0 |
| Recent BMI^*^, kg/m^2^ | |  |  |  |  |  |
|  | <25 | 3197 | 571 (24.7) | 600 (31.3) | 971 (45.5) | 1055 (51.4) |
|  | 25-30 | 3275 | 1113 (48.1) | 940 (49.1) | 631 (29.6) | 591 (28.8) |
|  | >30 | 1864 | 611 (26.4) | 372 (19.4) | 501 (23.5) | 380 (18.5) |
|  | Unknown | 76 | 17 (0.7) | 4 (0.2) | 30 (1.4) | 25 (1.2) |
| Cigarette smoking, pack-years | |  |  |  |  |  |
|  | Never | 3549 | 810 (35) | 689 (36) | 1003 (47) | 1047 (51) |
|  | <10 | 1368 | 332 (14.4) | 281 (14.7) | 392 (18.4) | 363 (17.7) |
|  | 10 to 19 | 928 | 289 (12.5) | 229 (12) | 236 (11.1) | 174 (8.5) |
|  | 20+ | 2238 | 807 (34.9) | 632 (33) | 411 (19.3) | 388 (18.9) |
|  | Unknown | 329 | 74 (3.2) | 85 (4.4) | 91 (4.3) | 79 (3.9) |
| Alcohol Consumption, No. of drinks/week | |  |  |  |  |  |
|  | 0 | 3421 | 703 (30.4) | 578 (30.2) | 1108 (51.9) | 1032 (50.3) |
|  | > 0, ≤7 | 2735 | 683 (29.5) | 654 (34.1) | 686 (32.2) | 712 (34.7) |
|  | >7, ≤14 | 999 | 362 (15.7) | 344 (18) | 151 (7.1) | 142 (6.9) |
|  | >14 | 944 | 470 (20.3) | 310 (16.2) | 87 (4.1) | 77 (3.8) |
|  | Unknown | 313 | 94 (4.1) | 30 (1.6) | 101 (4.7) | 88 (4.3) |
| Vegetable consumption, servings/day | |  |  |  |  |  |
|  | <2 | 4318 | 1412 (61.1) | 1212 (63.3) | 874 (41) | 820 (40) |
|  | 2+ | 4005 | 873 (37.8) | 683 (35.6) | 1228 (57.6) | 1221 (59.5) |
|  | Unknown | 89 | 27 (1.2) | 21 (1.1) | 31 (1.5) | 10 (0.5) |
| Fruit consumption, servings/day | |  |  |  |  |  |
|  | <1 | 2505 | 910 (39.4) | 643 (33.6) | 525 (24.6) | 427 (20.8) |
|  | 1+ | 5670 | 1330 (57.5) | 1216 (63.5) | 1536 (72.0) | 1588 (77.4) |
|  | Unknown | 237 | 72 (3.1) | 57 (3.0) | 72 (3.4) | 36 (1.8) |
| Red meat consumption, servings/day | |  |  |  |  |  |
|  | <1 | 6580 | 1681 (72.7) | 1484 (77.5) | 1721 (80.7) | 1694 (82.6) |
|  | 1+ | 1444 | 564 (24.4) | 364 (19.0) | 288 (13.5) | 228 (11.1) |
|  | Unknown | 388 | 67 (2.9) | 68 (3.5) | 124 (5.8) | 129 (6.3) |
| Physical activity (current vigorous exercise, hr/wk) | |  |  |  |  |  |
|  | 0 | 1583 | 469 (20.3) | 394 (20.6) | 383 (18.0) | 337 (16.4) |
|  | > 0, ≤2 | 3015 | 747 (32.3) | 681 (35.5) | 781 (36.6) | 806 (39.3) |
|  | >2, ≤4 | 1181 | 310 (13.4) | 317 (16.5) | 256 (12.0) | 298 (14.5) |
|  | >4 | 1421 | 447 (19.3) | 338 (17.6) | 351 (16.5) | 285 (13.9) |
|  | Unknown | 1212 | 339 (14.7) | 186 (9.7) | 362 (17) | 325 (15.8) |
| Calcium supplement use duration, years | |  |  |  |  |  |
|  | Non-user | 5923 | 2060 (89.1) | 1610 (84) | 1268 (59.4) | 985 (48.0) |
|  | ≤ 2.5 | 785 | 115 (5.0) | 97 (5.1) | 294 (13.8) | 279 (13.6) |
|  | > 2.5 | 1163 | 84 (3.6) | 137 (7.2) | 378 (17.7) | 564 (27.5) |
|  | Unknown | 541 | 53 (2.3) | 72 (3.8) | 193 (9.0) | 223 (10.9) |
| Regular NSAID use† duration, years | |  |  |  |  |  |
|  | Non-user | 4703 | 1364 (59.0) | 939 (49.0) | 1319 (61.8) | 1081 (52.7) |
|  | ≤ 2 | 1701 | 475 (20.5) | 388 (20.3) | 396 (18.6) | 442 (21.6) |
|  | > 2 | 1632 | 406 (17.6) | 509 (26.6) | 288 (13.5) | 429 (20.9) |
|  | Unknown | 376 | 67 (2.9) | 80 (4.2) | 130 (6.1) | 99 (4.8) |
| History of FOBT‡ | |  |  |  |  |  |
|  | No | 5447 | 1569 (67.9) | 1175 (61.3) | 1502 (70.4) | 1201 (58.6) |
|  | Yes | 2647 | 646 (27.9) | 667 (34.8) | 541 (25.4) | 793 (38.7) |
|  | Unknown | 318 | 97 (4.2) | 74 (3.9) | 90 (4.2) | 57 (2.8) |
| History of polyp‡ | |  |  |  |  |  |
|  | No | 7548 | 2079 (89.9) | 1691 (88.3) | 1943 (91.1) | 1835 (89.5) |
|  | Yes | 662 | 159 (6.9) | 197 (10.3) | 129 (6.0) | 177 (8.6) |
|  | Unknown | 202 | 74 (3.2) | 28 (1.5) | 61 (2.9) | 39 (1.9) |
| History of screening sigmoidoscopy‡ | |  |  |  |  |  |
|  | No | 6509 | 1893 (81.9) | 1397 (72.9) | 1747 (81.9) | 1472 (71.8) |
|  | Yes | 1547 | 330 (14.3) | 434 (22.7) | 281 (13.2) | 502 (24.5) |
|  | Unknown | 356 | 89 (3.8) | 85 (4.4) | 105 (4.9) | 77 (3.8) |
| History of screening colonoscopy‡ | |  |  |  |  |  |
|  | No | 7096 | 2035 (88.0) | 1536 (80.2) | 1890 (88.6) | 1635 (79.7) |
|  | Yes | 1119 | 217 (9.4) | 332 (17.3) | 197 (9.2) | 373 (18.2) |
|  | Unknown | 197 | 60 (2.6) | 48 (2.5) | 46 (2.2) | 43 (2.1) |
| Postmenopausal hormones use | |  |  |  |  |  |
|  | Non-user | 2538 | -- | -- | 1591 (74.6) | 1347 (65.7) |
|  | Estrogen only | 484 | -- | -- | 207 (9.7) | 277 (13.5) |
|  | Estrogen + Progesterone only | 309 | -- | -- | 126 (5.9) | 183 (8.9) |
|  | Mixed | 241 | -- | -- | 96 (4.5) | 145 (7.1) |
|  | Unknown | 4440 | -- | -- | 113 (5.3) | 99 (4.8) |

BMI: Body mass index; FRP: Familial Risk Profile; FH: binary family history; FOBT: fecal occult blood test; NSAID: Nonsteroidal anti-inflammatory drugs.

* As of two years before enrollment

† Regular NSAID use was defined as use of aspirin and/or ibuprofen at least twice a week for more than a month

‡ History of polyp, FOBT, sigmoidoscopy and colonoscopy were defined as history of having any of these conditions/test two years prior to enrollment.

**Supplementary Table 2. Percent missing for covariates in the validation set by sex***

| **Covariate** | **%** |
| --- | --- |
| Male |  |
| Body mass index | 3.7 |
| Red meat consumption | 4.8 |
| NSAID use duration | 3.5 |
| Calcium supplement use duration | 2.0 |
| Cigarette smoking, pack-years | 2.1 |
| History of FOBT | 3.5 |
| History of polyps | 1.6 |
| History of sigmoidoscopy | 2.4 |
| History of colonoscopy | 1.2 |
| Female |  |
| Body mass index | 4.5 |
| Fruit intake | 4.5 |
| Red meat consumption | 8.5 |
| Calcium supplement use duration | 7.7 |
| Cigarette smoking, pack-years | 2.9 |
| History of FOBT | 2.9 |
| History of polyps | 1.9 |
| History of sigmoidoscopy | 2.4 |
| History of colonoscopy | 1.4 |
| Hormone replacement therapy | 5.5 |

* FOBT: fecal occult blood test; NSAID: Nonsteroidal anti-inflammatory drugs.

**Supplementary Table 3. Comparison of model performance using age-adjusted AUC with and without imputation by sex*, by population vs clinic-based families**

| **Sex and model** | **Age-adjusted AUC (95% CI)** | |
| --- | --- | --- |
|  | **With Imputation** | **Without Imputation** |
| **Population-based** |  |  |
| Male FRP | 0.69 (0.60-0.78) | 0.70 (0.61-0.79) |
| Male FH | 0.61 (0.52-0.71) | 0.62 (0.53-0.71) |
| Female FRP | 0.70 (0.62-0.77) | 0.69 (0.61-0.77) |
| Female FH | 0.60 (0.52-0.67) | 0.58 (0.51-0.66) |
| **Clinic-based** | | |
| Male FRP | 0.77 (0.69-0.84) | 0.77 (0.70-0.84) |
| Male FH | 0.66 (0.58-0.74) | 0.66 (0.58-0.74) |
| Female FRP | 0.68 (0.60-0.76) | 0.68 (0.61-0.76) |
| Female FH | 0.57 (0.49-0.65) | 0.57 (0.49-0.65) |

*AUC = area under the curve; CI = confidence interval; FH = family history; FRP = familial risk profile

**Supplementary Figures**

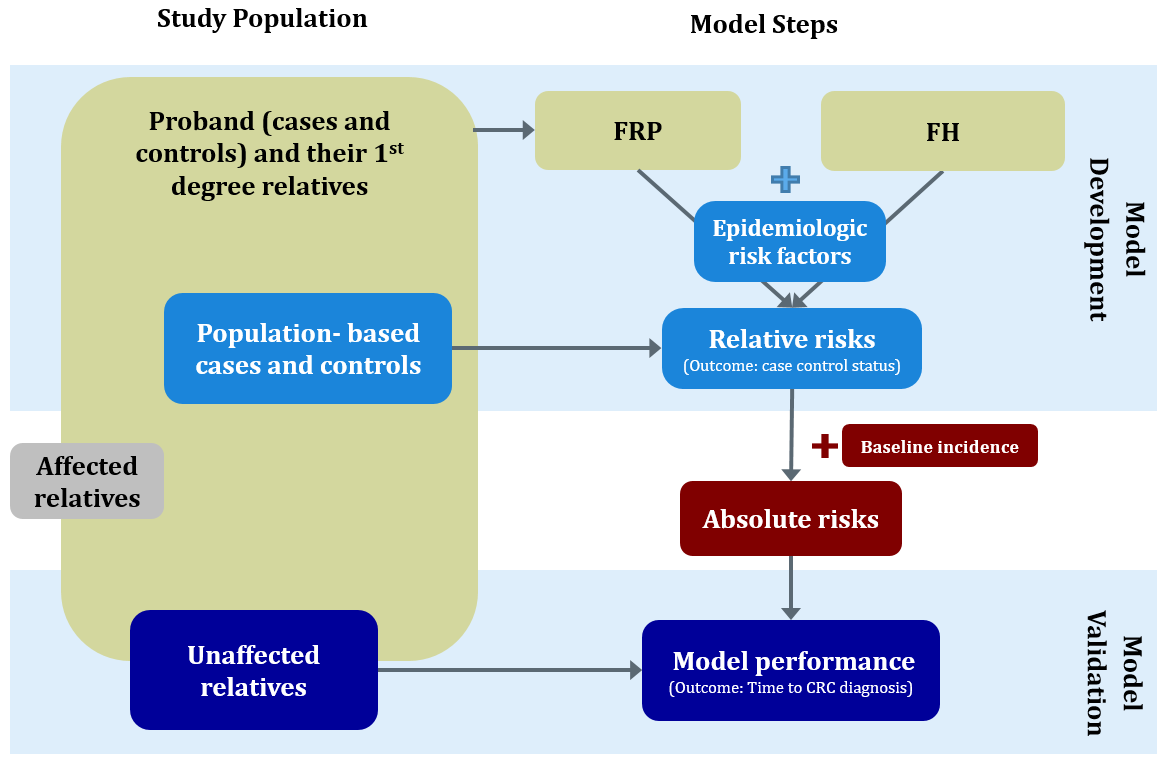

**Supplementary Figure 1. Study population and model steps. FRP: familial risk profile model; FH: binary family history model**

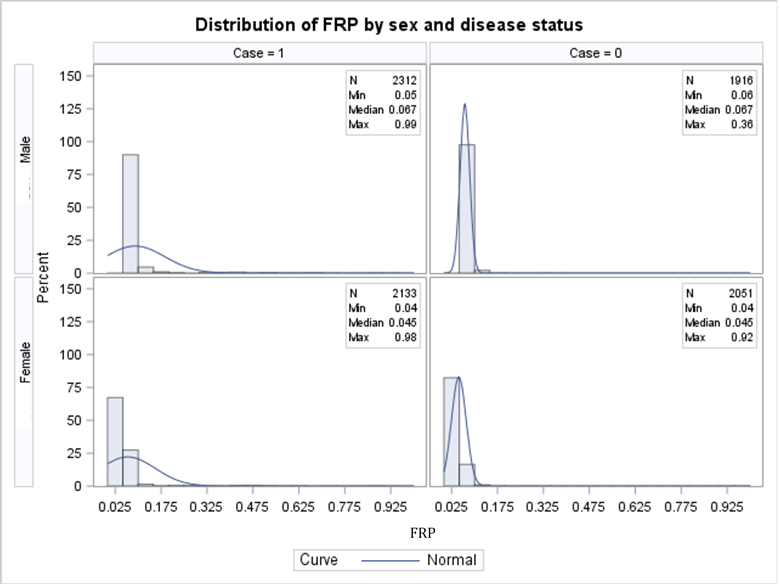

**Supplementary Figure 2. Histogram of the Familial Risk Profile (FRP) by sex and by case-control status**

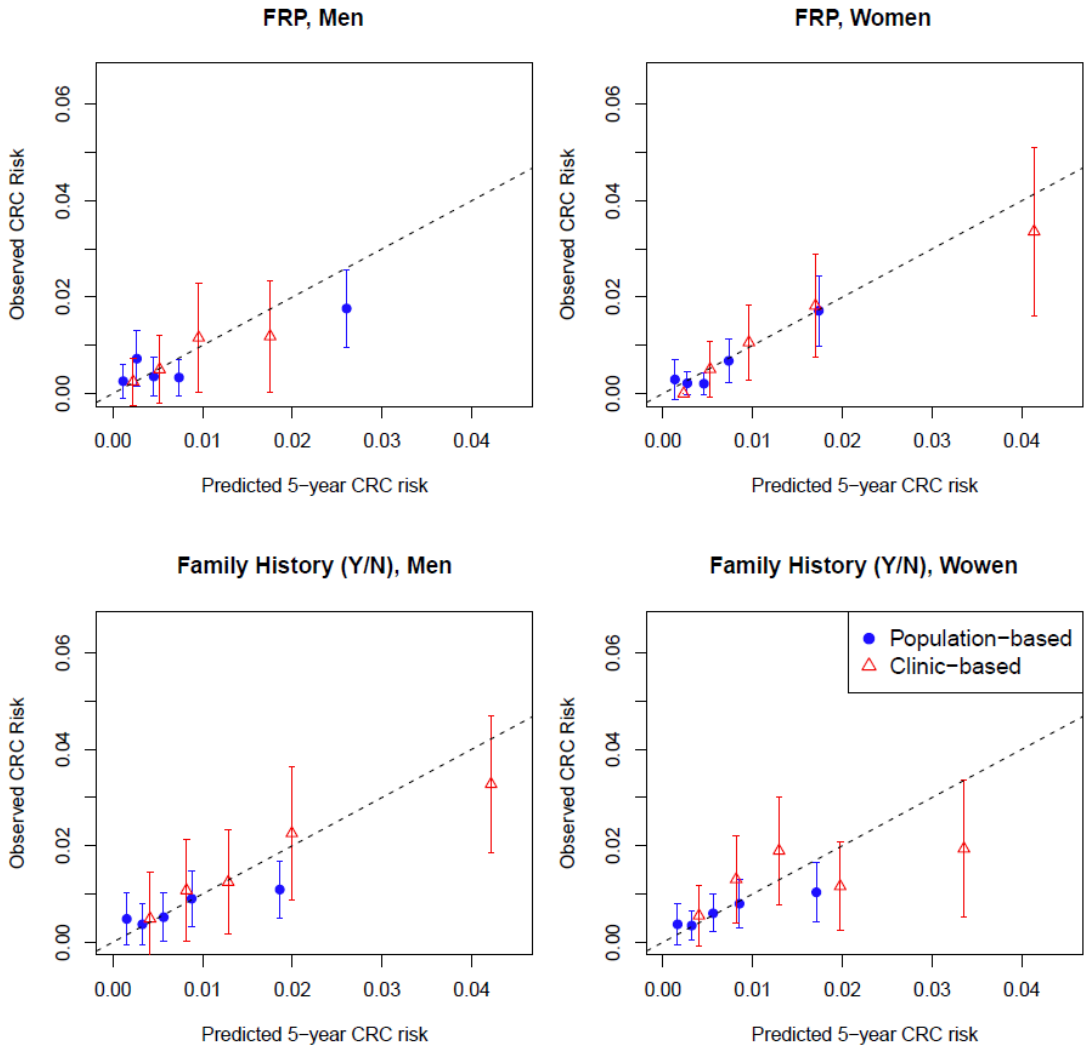
**Supplementary Figure 3.** Calibration plots by model and sex. Average observed 5-year colorectal cancer (CRC) risk (with 95% confidence intervals as error bars) vs averaged predicted 5-year CRC risk. Five groups were categorized by quintiles of predicted absolute risks. The observed risks and 95% CI were calculated as the cumulative incidences of CRC within 5 years accounting for censoring and competing risks. FRP: familial risk profile.
